# Random Forest Factorization Reveals Latent Structure in Single Cell RNA Sequencing Data

**DOI:** 10.1101/2021.09.13.460168

**Authors:** Boris M. Brenerman, Benjamin D. Shapiro, Michael C. Schatz, Alexis Battle

## Abstract

Single-cell RNA sequencing data contain patterns of correlation that are poorly captured by techniques that rely on linear estimation or assumptions of Gaussian behavior. We apply random forest regression to scRNAseq data from mouse brains, which identifies the co-regulation of genes within specific cellular contexts. By analyzing the estimators of the random forest, we identify several novel candidate gene regulatory networks and compare these networks in aged and young mice. We demonstrate that cell populations have cell-type specific phenotypes of aging that are not detected by other methods, including the collapse of differentiating oligodendrocytes but not precursors or mature oligodendrocytes.

## Background

Single-cell RNA sequencing (scRNAseq) is a powerful and widely-used technique to study transcriptome dynamics, especially within heterogeneous cell populations. However, scRNAseq data present challenges in interpretation due to sparsity, high dimension, non-Gaussian behavior, and other confounding effects (Hafemeister and Satija 2019). There are four major lines of inquiry that scRNAseq is commonly used to investigate: 1.) discovery and annotation of cell types, 2.) discovery and annotation of gene regulatory relationships, 3.) changes in cell type populations or gene expression across multiple samples, and 4.) the relationships of cell types to each other across differentiation trajectories and pseudotime (Papalexi and Satija 2018).

Principal component analysis (PCA) and related methods such as kernel PCA and projection pursuit regression (PPR) are widely used techniques to aid in the interpretation of scRNA data by consolidating redundant information across multiple genes into a more compact representation. First, PCA can be used to make operations such as clustering faster by allowing clustering methods to operate on lower dimensional data as used in e.g. Seurat (Hao et al. 2021). PCA is also often used to simplify the behavior of many genes into a single, simple explanatory variable that can be interpreted to better understand a dataset. For example, if the first principal component (PC1) is effective in separating samples that underwent a stimulus from control samples, an experimenter might infer that the weights of PC1 will summarize the response of samples to the stimulus. (Jolliffe and Cadima 2016)

However, this use of PCA can be dangerous because PCA makes the assumption that the relationships between the variables (genes) in the data are consistent across all samples observed and the features have only strictly linear relationships. It also assumes the data can be meaningfully centered because the framework of PCA relies on projecting data into a subspace defined by the variance of the input data around an arithmetic mean. In practice, gene expression data for scRNAseq is often highly skewed and contains non-linear dependencies between genes even after transformation. Such nonlinear dependencies between features are often referred to as “higher order dependencies” and violate the basic assumption of PCA (Hyvärinen 1999). For example, transcriptional response to DNA damage is different in cycling cells, which prefer homologous recombination as a response to double-stranded breaks and non-cycling cells, which prefer non-homologous end-joining. The practical outcome of analyzing data with non-linear dependencies by PCA is that PCs will conflate both the relationships between groups of genes that in reality operate independently, or are even anti-correlated with each other. For example, a PC that captures DNA damage response may well weigh both NHEJ and HR highly, implying they are equally preferred in all cells that experience DNA damage, while in practice one is usually chosen over the other. (Iyama and Wilson 2013)

Kernel PCA and Projection Pursuit Regression (PPR) can both be used to address this problem (Townes et al. 2020), but kernel PCA does not provide any meaningfully interpretable feature weights, and PPR can be quite slow and is sensitive to the selection of a good objective. Certain aspects of non-linear behavior can be overcome by performing clustering and analysis in embedded space using tSNE or UMAP, but this practice has fallen out of favor by major packages (eg scanpy, Seurat, and Monocle no longer cluster in embedded space though they have in the past) because of the difficulty in interpreting the relationship between the embedding and the source data; for example reverse transformation out of a tSNE embedding is impossible. Regardless, clustering in embedded space does not provide one with insight regarding the relationships between correlated variables, so it can’t be understood as a conventional factoring method. Non-negative matrix factorization is also sometimes called for in interpreting cell signatures (as in the ProjectR package) but has similar limitations as tSNE embedding, such as the difficulty in interpreting the significance of any particular signature. In practice, the field is in need of a method that will allow us to find simple and intuitive factor decompositions of single-cell RNAseq data that both allow for improved clustering in a lower-dimensional space and simultaneously provide intuitive interpretations for the redundant information spread across many genes. In addition, the method should enable the interpretation of how various regulatory relationships interact with each other in a complex landscape and how they vary across the dataset. Ideally, we would want to find a method that identifies broad divisions present among cell types (e.g. stem, differentiated, cycling, quiescent, etc.), as well as the gene regulatory relationships present within those divisions.

To address the problems posed by higher-order dependencies, we consider the use of random forest regression (RFR). Random forests (RF) are a class of statistical models that are ensemble estimators based on bootstrapped decision trees. They are non-linear, make few assumptions about the distribution of the underlying data, are robust in a variety of contexts, and are accurate and efficient (Breiman 2001). Unfortunately, RFs are not straightforward to interpret if a user seeks to understand the behavior of correlated features or obtain a meaningful high-level representation of a dataset, such as that provided by principal components or cluster assignments. The availability of measures such as feature importance offer insight into individual features but not the coordinated behavior of several features in the data.

In order to more effectively understand the structure of datasets such as single-cell gene expression measurements, we propose a novel method for the analysis of the latent structure of an RF. To obtain a general characterization of the data with few initial assumptions, we propose a technique for training the RFR in an unsupervised manner on a gene expression matrix, rather than regressing on a single output variable, allowing the RFR to predict the behavior of all genes based on the expression of all other genes. To understand the decisions that RFs make to predict gene behavior, we propose examining the commonalities across the individual nodes making up an RF directly. Given RF nodes that have many properties in common, we can cluster them and infer what we call *random forest factors* (RFFs). Random forest factors trained on scRNA-seq data partition cells into groups, capture the gene covariances present within each group, and covariances present across several groups. RFFs have simple and intuitive criteria for group membership that can be extrapolated to other datasets (**Figure 1**). This approach provides a way to naturally capture higher-order interactions in single cell gene expression data, provide a partitioning scheme that is suited to irregularly-shaped clusters, produce interpretable summaries of many features, and enable comparisons between distinct datasets. When applied to mouse brain scRNAseq data, we are able to pinpoint significant changes in gene expression for differentiating oligodendrocytes as a function of aging, aiding in the interpretation of otherwise opaque data.

**Figure 1.**
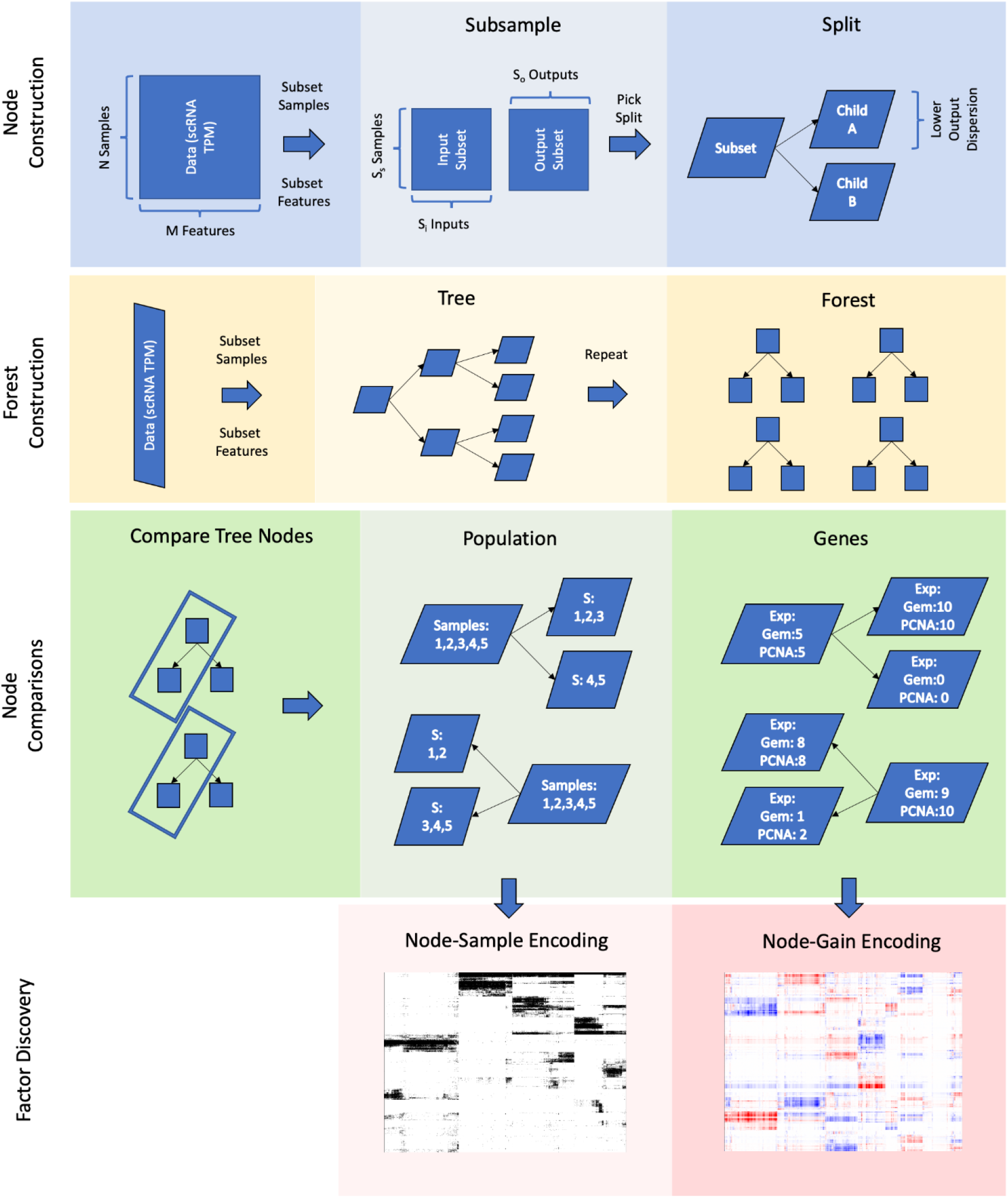
Overview of Random Forest construction. Node construction: to construct a node, the raw data matrix is bootstrapped by sample with replacement (eg rows are randomly selected and copied into a new matrix), then the sample bootstrap matrix columns are divided into inputs and outputs without replacement, (eg the order of the columns is randomized and the matrix is split in two). The input and output columns are then bootstrapped with replacement producing the node input and node output matrices of a specified dimension. The node is split by evaluating different splitting criteria. To construct a forest we repeat the node construction procedure several times using different bootstraps to create a collection of different decision trees. In turn to interpret nodes we examine the samples present in each node and the regression a node performs on the samples it contains when it is split into children.

## Results

### Methodology

To understand scRNAseq data better, we must have ways of identifying cell populations with common expression profiles, identifying which expression profiles distinguish a cell population, identifying how the genes interact with each other, and identifying which genes are co-regulated to determine the traits of a cell. In a context such as scRNAseq where there are higher-order interactions, we can also ask whether or not genes and traits behave in the same way across all cell populations, or whether or not their behavior changes in some cell populations.

An RFR is an ensemble estimator based on a collection of individual decision tree estimators, which are in turn collections of individual nodes. In order to generate a tree, each individual node partitions a set of samples into two subsets based on gene expression or another criterion and generates two child nodes corresponding to the subsets. For example, a root node might coarsely partition all cells in an scRNAseq experiment into cycling and non-cycling cells based on the expression level of Geminin, a DNA replication inhibitor (McGarry and Kirschner 1998) One subset would then contain only cells that express high levels of Geminin, and the other would contain only cells expressing low levels of Geminin; cycling and non-cycling cell subsets will then have lower gene expression variance than the original set of all cells. After a node is created and a criterion is selected that determines whether or not a sample should be partitioned by that node, that node becomes an *estimator* and a *partition*. Henceforth, in order to estimate the behavior of a new sample, we find if it satisfies the criteria of a node (e.g. its Geminin expression is high) and that node then predicts that the sample will be similar to the samples in its subset. A node that uses Geminin as a criterion might predict that samples partitioned into it will also have high expression of the polymerases, both for DNA replication and repair.

To analyze the behavior of a complex dataset such as scRNAseq expression data we will use a random forest regressor with multiple outputs constructed in an unsupervised manner. We construct the RFR with no supervision by iteratively and randomly splitting all genes into two subsets of inputs and outputs, and training trees to predict the expression of a set of output genes from the expression values of a set of input genes. This approach allows us to construct a random forest that relates the expression of each gene to the expression of other genes using bootstrapping, since we have no need to predict any known ground truth about the dataset.

RF nodes have an essential property that distinguishes them from other estimators: they are recursively generated. An RF node generates child nodes by finding the optimal split of the cells in its subset, without taking into account any other splits and based on entirely new criteria. In this way, each node makes a split on a population of cells where the variance explained by its parent node has been eliminated conditional on some criterion, which makes it a *conditional estimator*, otherwise known as a marginal estimator. Because each conditional estimator contains samples that meet criteria such as Geminin expression, the effects discovered by conditional estimators could be dependent on the levels of expression of previously used genes. For example, repair of double-stranded breaks can proceed either by non-homologous end-joining (NHEJ) or homologous recombination. Homologous recombination is more prevalent in cells that are cycling while NHEJ is more common in cells that are quiescent. Within an RF, an individual node might contain only cycling cells, or only quiescent cells, as determined by Geminin levels. Such a node can detect that expression of HR pathway genes is dependent on high Rad51 and Geminin expression simultaneously. However, if Geminin expression is low, Rad51 expression would have a small but positive correlation with NHEJ (Iyama and Wilson 2013). The fact that Rad51 predicts HR in cycling cells but NHEJ in quiescent cells is an example of a higher-order interaction. In this way conditional estimators allow for the possibility of discovering higher-order interactions such as conditional gene expression, which distinguishes them from purely linear estimators such as PCA.

### Methodology Overview

To compare nodes across different trees in a random forest, we will describe them in terms of the samples they contain (creating a node-sample encoding or NSE) or the improvement in the conditional prediction they make (creating a node-gain encoding or NGE). By comparing the conditional estimates a node makes and samples in that node, we will be able to judge the similarities of two nodes to each other, allowing us to determine if nodes can be arranged into meaningfully separated clusters via hierarchical or community clustering. Then, by examining nodes that cluster together, we will be able to determine other properties of those nodes such as conditional covariance of genes or conditional predictive power.

In order to analyze whether or not similar *conditional estimates* are frequently discovered in an RFR, we construct a summary of the predictions that a node makes relative to its parent. A ***node-gain encoding (NGE)*** is a matrix of size N x L where N is the number of nodes and L is the number of regression targets (gene expression values), with each element representing a regression target; an element is valued as the difference in the predicted value for a target in a given node compared to its parent, hereafter known as ***conditional gain (EQ1)***.

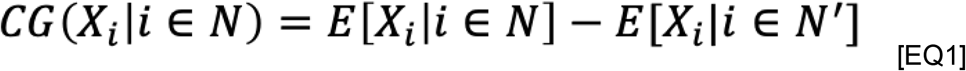

Where CG is conditional gain, X_i_ is the regression target X for sample i, N is a node of interest and N’ is the parent of the node of interest. In the case of the root node, which has no parent, the expectation of X given the parent is 0, so the conditional gain is the mean across the dataset. Consequently, conditional gain is the gain after controlling for all previously explained variation.

A **partial gain encoding** (PGE) is similar to a node-gain encoding but controls the change in the mean value for a particular node for the effects predicted by both its parent nodes and child nodes using the law of total variance (EQ2).

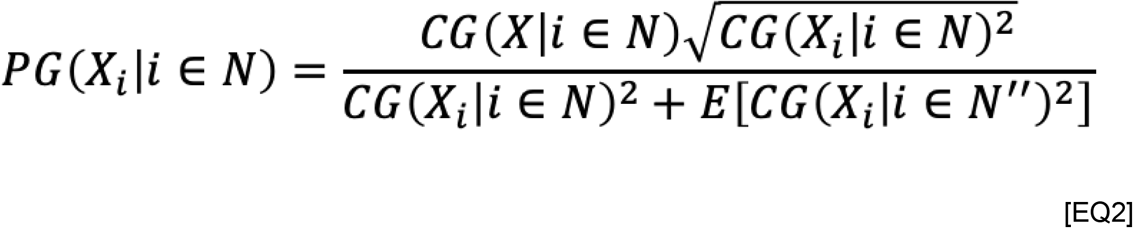

where PG is partial gain, CG is conditional gain, X_i_ is regression target X for sample i, N is a node of interest and N’’ any descendant of the node of interest or its sisters. In practice, we get a value between -1 and 1 whose magnitude is the percentage of total variance explained for a particular gene by node N. This measure is useful because the behavior of a given sample which is partially explained by the conditional gain of a given node N may in fact be explained by a node dependent on N (i.e. one of its children). Thus when examining the effects that a given node predicts most accurately, we want to eliminate predictions that another node may make more accurately still. By using the law of total variance, we scale the explanatory power of every node to the same scale while retaining the sign of the estimate.

The NGE and PGE allow us to summarize the conditional gain that node is predicting, so by comparing the conditional effects predicted by different nodes we can check whether certain predictions happen repeatedly. We are interested in examining these encodings because they allow us to see if nodes predict conditional changes in gene expression for the same genes repeatedly, implying that those genes are co-regulated. For example, if we find that many nodes predict that both Geminin and polymerase epsilon are upregulated simultaneously, our conclusion would be that there is a regulatory relationship between these two genes. However, the advantage compared to conventional correlation analysis like WGCNA is that many of the correlations we are examining are greatly controlled for correlations addressed by other estimators.

In order to compare the samples partitioned into nodes, we can construct a **node-sample encoding (NSE)**. A node-sample encoding is a matrix of size N x S, where N is the number of all nodes in an RFR and S is the number of samples; element (i,j) is 1 if node i contains element j and 0 otherwise. We constructed node sample encodings in order to examine whether certain groups of nodes often contained similar groups of samples. A node-sample score is the mean of the NSE for a given set of nodes, representing the proportion of nodes in a cluster of nodes that observe a certain sample.

A ***node-sister encoding*** is similar to a node-sample encoding, but can be valued as -1,0, or 1. An element in a node-sister encoding is valued 1 if a sample is partitioned into a node, -1 if it is partitioned into the sister of the node, and 0 if it is partitioned into neither. A node-sister encoding contains more information than a node-sample encoding, and allows us to visualize whether we we repeatedly see a single subset partitioned away from different sets, or if we see the same greater set partitioned into the same two subsets repeatedly across different trees.

The node sister encoding allows us to compute the sample sister score for a given cluster, which is the number of times a particular sample was encountered in the nodes of a node cluster - the number of times that sample was encountered in their sister nodes. The advantage of the sister score is that it is close to 0 when there is no information about a sample or if the information is ambiguous (eg a sample is equally frequently discovered in a particular node or its sisters), but the score value is high when a sample is frequently discovered only in the node or only in the sisters.

### Single Cell Analysis

In order to apply our factoring procedure to single cell data we trained an RFR on scRNAseq data derived from disaggregated brains of 8 young mice, with an average of 2000 cells per brain, by Ximerakis et al. (Ximerakis et al. 2019) We recalculated the cell by gene expression matrix with an identical preprocessing pipeline as Ximerakis and obtained 16027 single cell expression profiles of disaggregated brain tissue belonging to 8 young mice. An RFR was trained in an unsupervised manner on the cells originating from young mice, trimmed, and node-sample and node-gain encodings were derived from all nodes remaining in the forest. (**Figure 2**)

**Figure 2.**
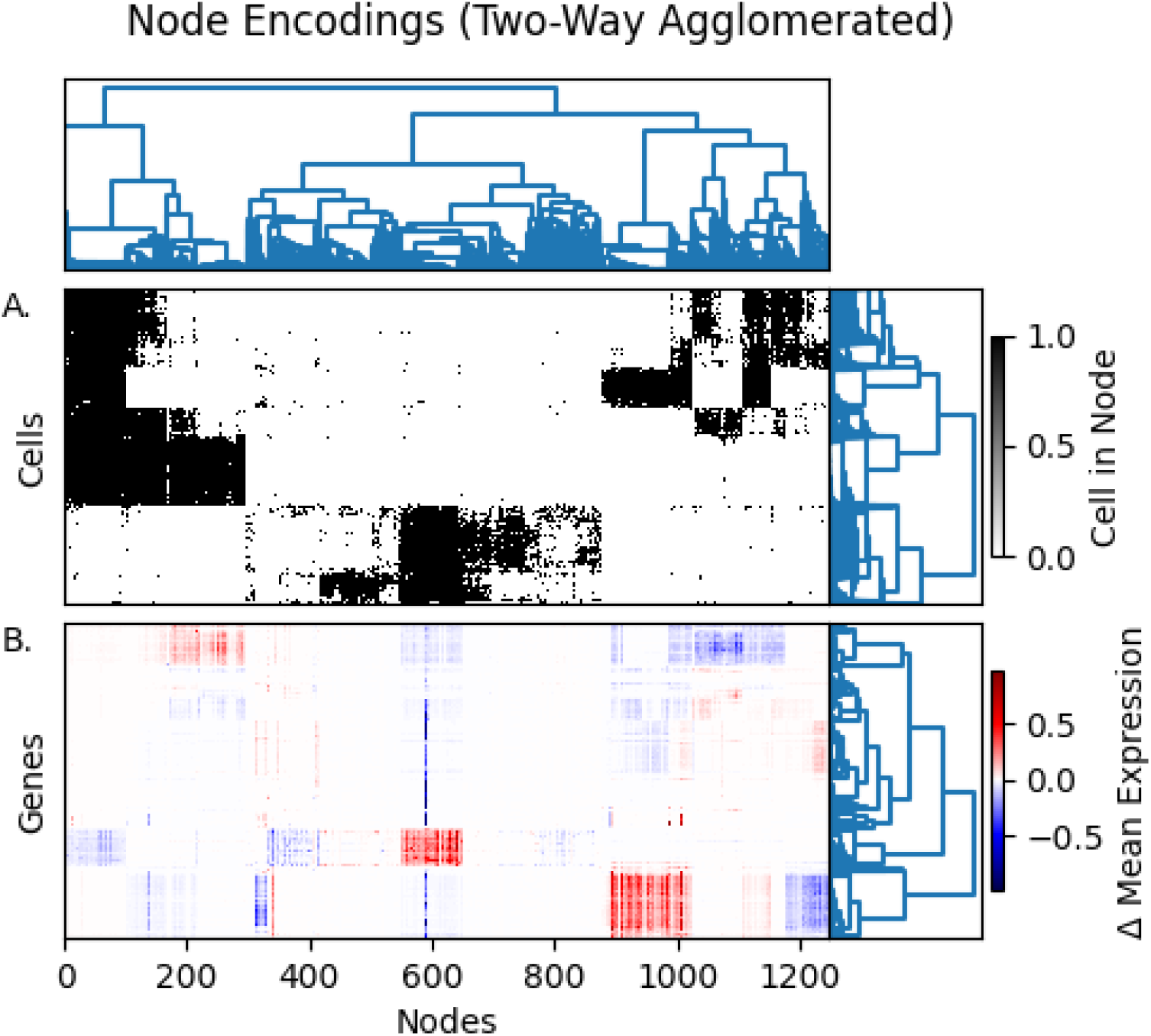
Node-Sample Encoding and Partial-Gain Encoding derived from an RFR trained on mouse single-cell gene expression data. Nodes below depth 4 are omitted for clarity, see Supplemental Figure 1 for complete figure. A) Node-sample encoding, element i,j is black if cell i is in node j, white otherwise. B) Partial-gain encoding, element i,j is red if expression of gene i is predicted to be higher by node j controlling for all other nodes in the tree, white if the expression is on average the same, and blue if the expression is predicted to be lower. Dendrograms are constructed by cosine similarity on a dimensionally reduced representation of data.

We agglomeratively linked both nodes and genes within the NGE using cosine similarity and observed a checker pattern indicating that nodes in the RFR repeatedly predict one of several categories of conditional gains (**Figure 2B**). The node subpopulations didn’t appear consistent in population or intra-cluster distance metrics so to obtain hard partitions rather than a dendrogram we computed a k-nearest-neighbors representation of the nodes and partitioned them via the Louvain algorithm into 39 clusters. 42% of all variance in the NGE was explained by the clustering procedure, reflecting the difficult nature of the underlying clustering task. We created several permutations of the NGE by shuffling each column independently to decorrelate the features. When the resulting permuted data was clustered by the same procedure, the resulting partitions explained 7.5 +/- 2e-7 % of the variance, indicating that the clusters found in the NGE correspond to real structure.

Importantly, when nodes are partitioned into clusters on the basis of the similarity of their NGE, the partitions explain 25% of the variance in the NSE as well, while partitioning an NSE that was shuffled as previously described explains only 7.3% of variance. Additional validation of the clustering performance has been performed on the iris dataset and is presented in supplemental figure 1.

Ultimately the usefulness of these clusters of nodes is in aggregating the effects of individual nodes into the mean effect across the cluster. This is necessary for interpreting the behavior of a random forest because bootstrapping prevents us from drawing confident conclusions about the behavior of any individual node. Whereas the behavior of a single node could be attributed only to chance inherent in the bootstrapping procedure, by taking a collection of nodes behaving in a similar manner, we can assert the presence of a particular effect.

### Local Behavior

The conditional dependencies demonstrated by the nodes in clusters can be different than the dependencies that are apparent across the dataset as a whole. For example, in the brain dataset RFR the behavior of immune cells is described by nodes falling into cluster 5 and its descendants, as evident from the gene expression of cells in those clusters. There is a clear trimodal distribution of cluster 5 sister scores (**Fig. 3A**), therefore we can compare cells that occur unambiguously in cluster 5 (>0.2 sister score) against all other cells. Samples that unambiguously occur in node cluster 5 significantly overexpress CD45 (.37 vs .016 TPM, p=1e-106, T Test) compared to all other cells, a ubiquitous immune cell surface marker (Trowbridge, Ralph, and Bevan 1975), and the C1Q family of genes (4.15 vs .04 TPM, p<machine epsilon, T Test), which are components of the complement system (Reid, Lowe, and Porter 1972), so cluster 5 broadly specifies immune cells. Nodes in Cluster 5 also predict the overexpression of C1Qa compared to parent nodes. The difference in mean expression of C1Qa in nodes of NC5 compared to their parents is 1.0. This is the *conditional gain* of C1qa in NC5.

**Figure 3:**
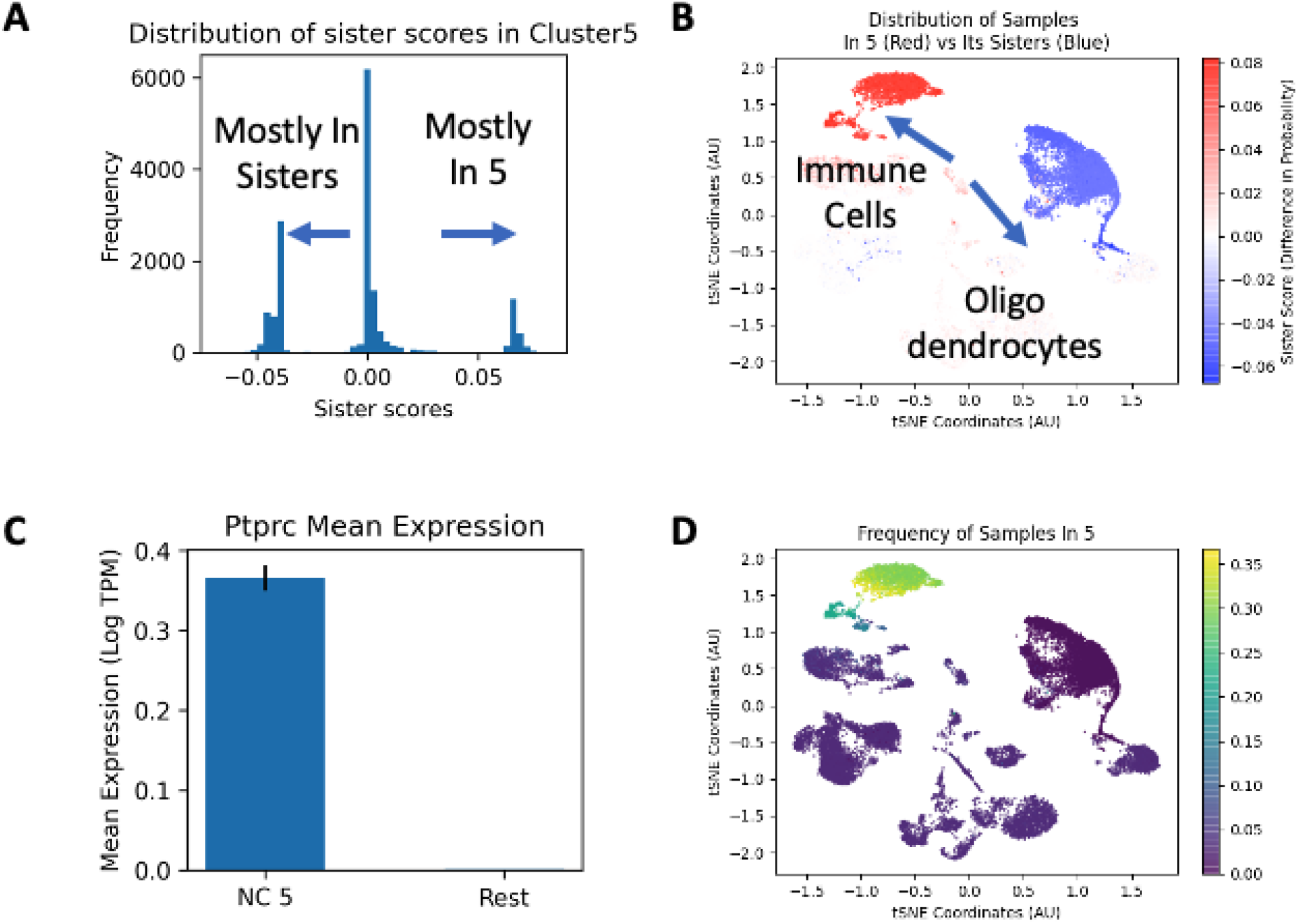
A. RFR nodes trained on mouse scRNAseq data and clustered by conditional gain have narrow distributions of sister scores. Samples occur mostly in nodes of cluster 5, mostly in its sisters, or mostly in neither. B. Samples that occur mostly in node cluster 5 are immune cells, as established by CD45 expression. C.Mean expression of Ptprc (CD45) in samples mostly occurring in NC5 and all other samples, whiskers are SEM. D. Frequency of samples in NC5 overall. Some samples occur frequently in both NC5 and its sisters and have a low sister score despite being seen frequently in NC5.

The latent variables that govern gene expression in mouse brain cells are not fully known, so we examined whether forest factors can discover local behavior by comparing the correlations discovered by forest factors to correlations captured by PCs and conventional correlations. Similar analysis has been performed on simulated data with nested structure and known latent variables, for further details please see supplemental figure 2.

The mouse brain samples we are examining contain both microglia and immune cells normally traveling in blood, likely as a consequence of blood vessels being present in the brain samples collected. Microglia precursors share a lineage with other immune cells, however they migrate into the brain during early stages of mouse embryonic development and thus the gene expression profiles of general immune cells and microglia can be grouped naturally. There is a very significant distinction between microglia and other immune cells as well, which may be of interest to us; neuronal-lineage cells have a drastically different gene expression profile from any immune-lineage cells, and the gene-regulatory programs that govern immune cells and neuronal cells have little overlap. Among immune cells which are described by node cluster 5 (NC5), the RFR detects the differential regulation of several key genes that distinguish microglia and macrophages, such as CD74, Tmem119, Lyz2, and H2-Ab1.

Nodes from NC5 make up 57% of parents of NC34 and 32% of the parents of NC27. Nodes in NC34 predict the over-expression of Tmem119, which is a marker of microglia (conditional gain .91). Conversely, Cluster 27 predicts the over-expression of CD74 (conditional gain .40) and Mrc1 (conditional gain .88), which are strong markers of perivascular macrophage identity in combination (Kim et al. 1992). Of the top 5 genes most over-expressed in NC34 compared to its sisters, all are strongly anti-correlated with the top 5 most over-expressed genes in NC27 relative to its sisters when the correlation is weighted by the frequency with which each cell is present in either of the two clusters (**Fig 4H**). However, globally the genes in these two subsets are uncorrelated or even positively correlated (**Fig 4G**). For example the expression of Tmem119 and Cd74 is uncorrelated globally, however, when examining only the behavior of cells frequently present in Cluster 34 but not its sisters, we observe a strong positive correlation, which is an example of *local behavior*. Part of the reason that this correlation may be difficult to observe in other ways lies in the cells belonging to node cluster 18, where Tmem119 and Cd74 are positively correlated (**Fig 4F**). NC18 represents arachnoid barrier cells, as established by expression of Slc47a1 relative to other vascular cells.

**Figure 4:**
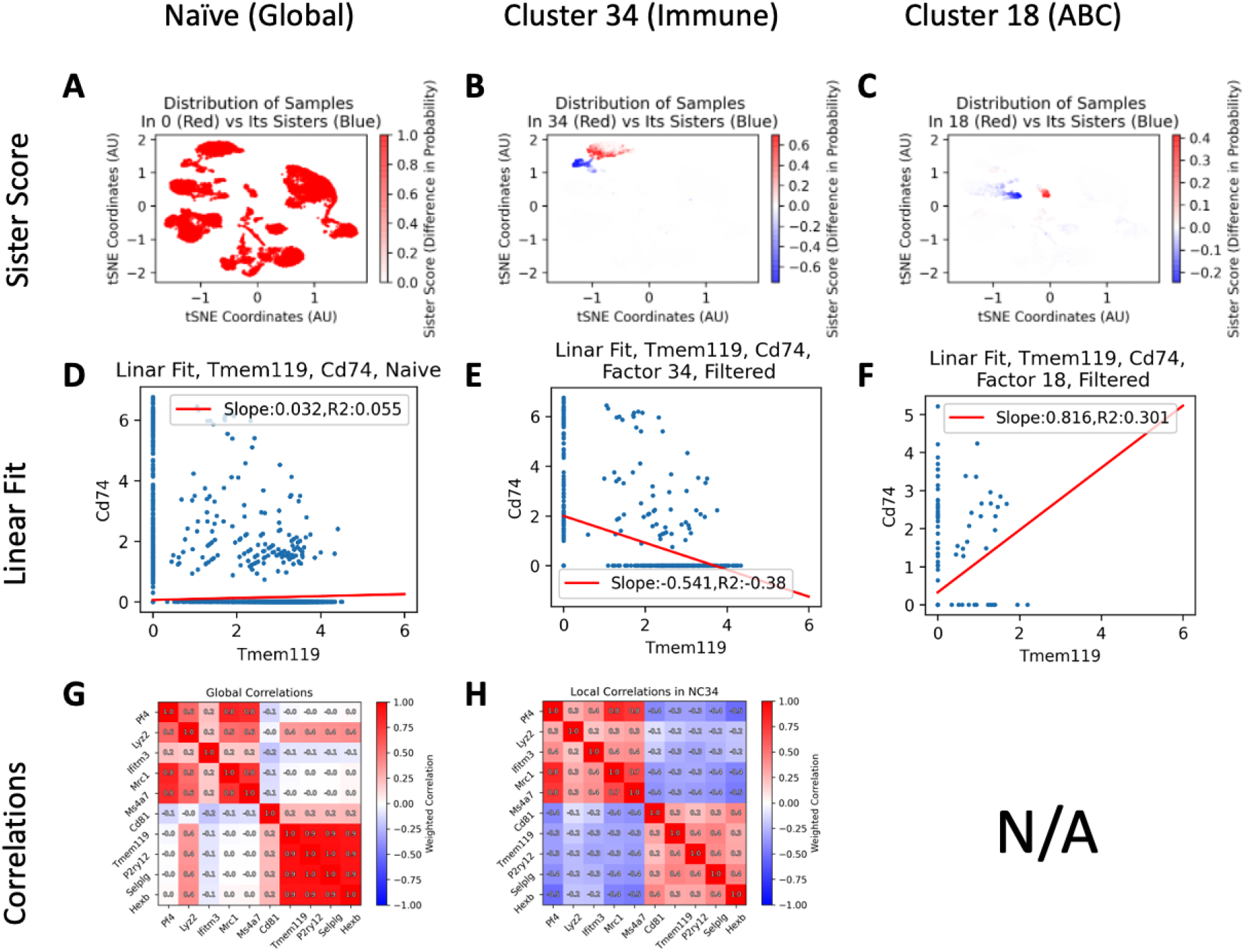
Certain genes behave differently when considering only cells frequently seen in NC34 than they do when considering all cells. **A**. UMAP projection of cell gene expression data. **B**. UMAP projection of cell gene expression data colored by random forest factor score for RFF 34. All cells express CD45 and are therefore immune, red cells express Tmem119 and are therefore microglia. **C**. UMAP projection of cell gene expression data colored by RFF 18. Red cells express Slc47a and are therefore arachnoid barrier cells.

Many genes have local behavior similar to CD74 and Tmem119, such as Pf4, Lyz2, Ifitm3, Mrc1, Ms4a7 and separately CD81, Tmem119, P2ry12, Selplg, and Hexb. Both these sets of genes are strongly anti-correlated with genes of the other set, but only among the cells unique to NC 34. (**Fig 4G, 4H**), and we propose that they occupy a gene regulatory block distinctive to microglia. Our ability to identify these sets of genes as being co-regulated in specific ways in different contexts is desirable since it allows us to identify a gene regulatory block that distinguishes two populations of a related lineage from each other, representing only the conditional difference between microglia and blood immune cells while controlling for irrelevant variation present in the rest of the dataset.

### Comparison to PCA

Finding this particular regulatory block using PCA is challenging, because each PC represents variation from several sources. When the log-normalized and scaled expression matrix is analyzed by PCA, the weights of individual genes in each component correspond to the regulatory relationships that PC is modeling. In order for us to find the regulatory program that distinguishes microglia from other immune cells we would need to find a PC which weighs Tmem119 and CD74 with opposite signs. We could then examine the weights of other genes in that PC and surmise that ones with weights of the same sign as Tmem119 and CD74 are upregulated in microglia and blood immune cells respectively.

**Supplemental:**
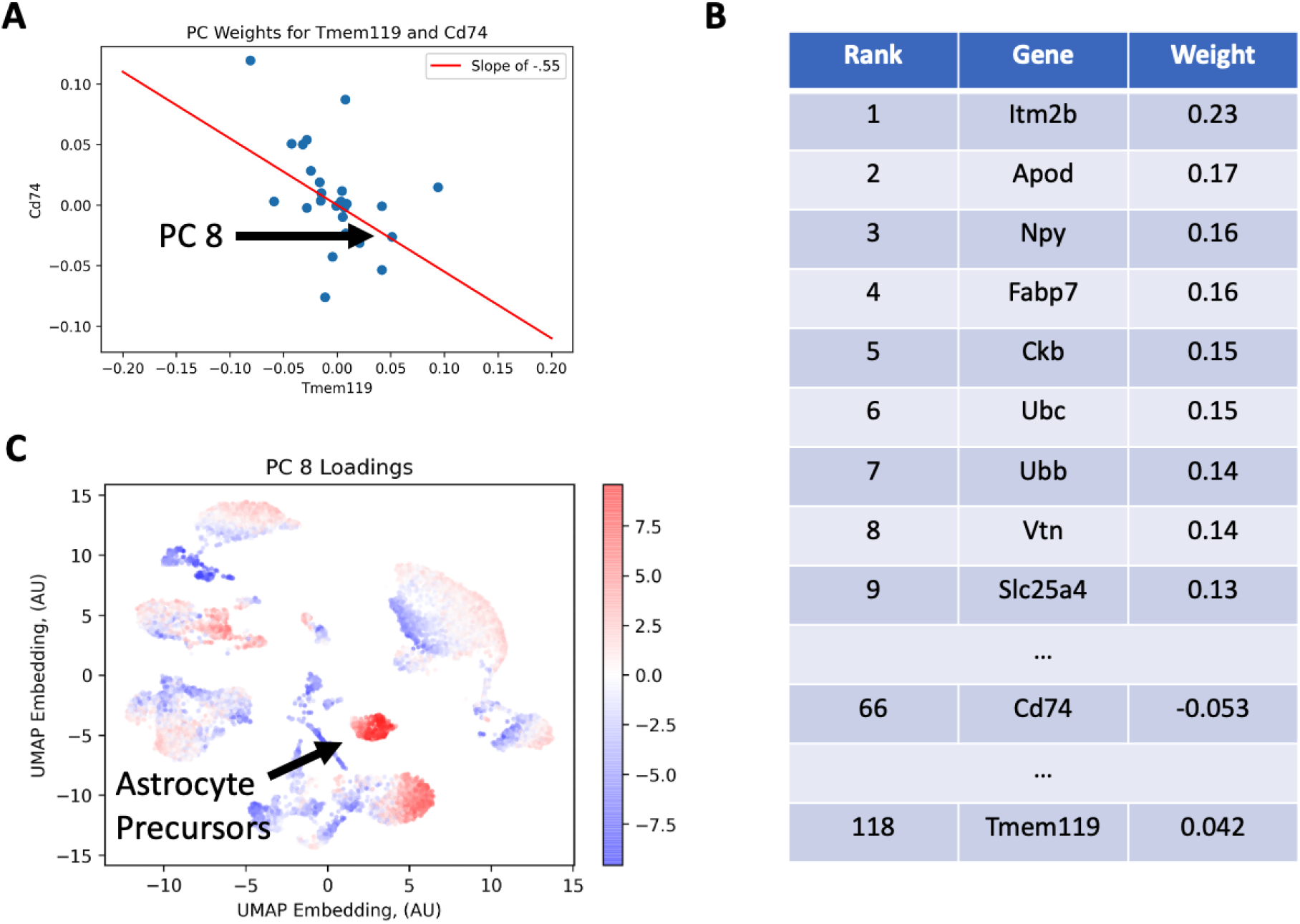
Plot of weights of Cd74 and Tmem119 among all PCs trained on mouse brain data. Red line has a slope of -.55, which is the same as the correlation coefficient of CD74 and Tmem119 among immune cells. For a PC to weigh Tmem119 and CD74 identically, it must lie on the red line. B: Genes ranked by the absolute value of their weighing in PC8 of the mouse brain dataset. C. Loadings of PC8 trained on mouse brain cells overlaid on a UMAP projection of all cells, red indicating a large positive loading, blue indicating a large negative loading. Endothelial cell identity established by

PC8 has weights of Tmem119: 0.042 and CD74:-0.053, which is the strongest negative association of loadings observed among the PCs and accurately captures the anti-correlation that is also captured by Cluster 34. Thus, it could be that the weights in PC8 model the distinction between microglia and macrophages. However, PC8 has loadings across a range of cell types, and high weights for genes that are unrelated to the behavior of the immune system. The gene with the second-highest weighing in PC8 is Apod, which is not known to be expressed in any immune cells (Yoshida et al. 1996), and thus unlikely to be co-regulated with Tmem119 and Cd74. Thus, if we were to use this PC to model the distinction between microglia and macrophages, we would find a number of genes in our model whose behavior is not relevant.

### Node Clusters Allow For Comparisons of Datasets

We’ve shown that Random Forest Factors can summarize the behavior of several features that are co-regulated in particular contexts such as specific cell types. RFFs can also be used to demonstrate that particular conditional effects change between different datasets. There are three ways for the expression of genes in a single-cell RNAseq sample to shift across two different sets of conditions: (1) due to changes in the proportions of particular cell types in the sample; (2) due to changes in the actual gene expression profile of cells within a cell type; or (3) due to technical differences between the two datasets.

Random Forests don’t present any unique advantage in compensating for technical artifacts, so proper normalization is required before they are applied as is the case for other methods, and we will assume it for subsequent analyses. If an RFR is used to partition a dataset in which identically behaving cells are present in different proportions, the accuracy of predictions for any particular node will not change; there will only be a change in the proportion of cells present in a particular set of nodes. Conversely, if gene expression within some cell types does change, then the model we have built to describe them is no longer valid. In this case, the quality of the predictions made by individual estimators in an RFR will change (presumably for the worse), since cells will be assigned to nodes that don’t fully describe their behavior. We can take advantage of this property in order to examine changes in populations and gene expressions between aged and young mouse brain cells. We used an RFR trained on young mouse brain cells to predict the behavior of aged mouse brain cells, which assigned aged brain cells to particular nodes in the RFR.

NC7 is a set of nodes describing the behavior of immature but committed oligodendrocytes, established by the differential expression of Cldn11 (oligodendrocyte transmembrane protein, 3.56 tpm in 7 vs 1.21 in all other^1^,p=4e-101) and Bmp4 (early differentiation factor, 0.424 tpm in 7 vs 0.015 in all other, p=1e-18).(Gow et al. 1999) To establish how well the nodes in NC7 can predict the behavior of immature oligodendrocytes we used bootstrapping. We partitioned the 8 mice from which brain samples were obtained into 35 possible partitions of 4 training mice and 4 test mice. (**Fig 5A**) When the cells from 4 training mice are used to establish the conditional effects of nodes in NC7 and predict the behavior of cells belonging to 4 test mice, we observe a mean COD of 11% and a range of CODs from 7.7% to 13%. (**Fig 5C**), indicating that 11% of the variance present among all oligodendrocytes can be explained by the unique gene expression profile of immature oligodendrocytes.

**Figure 5:**
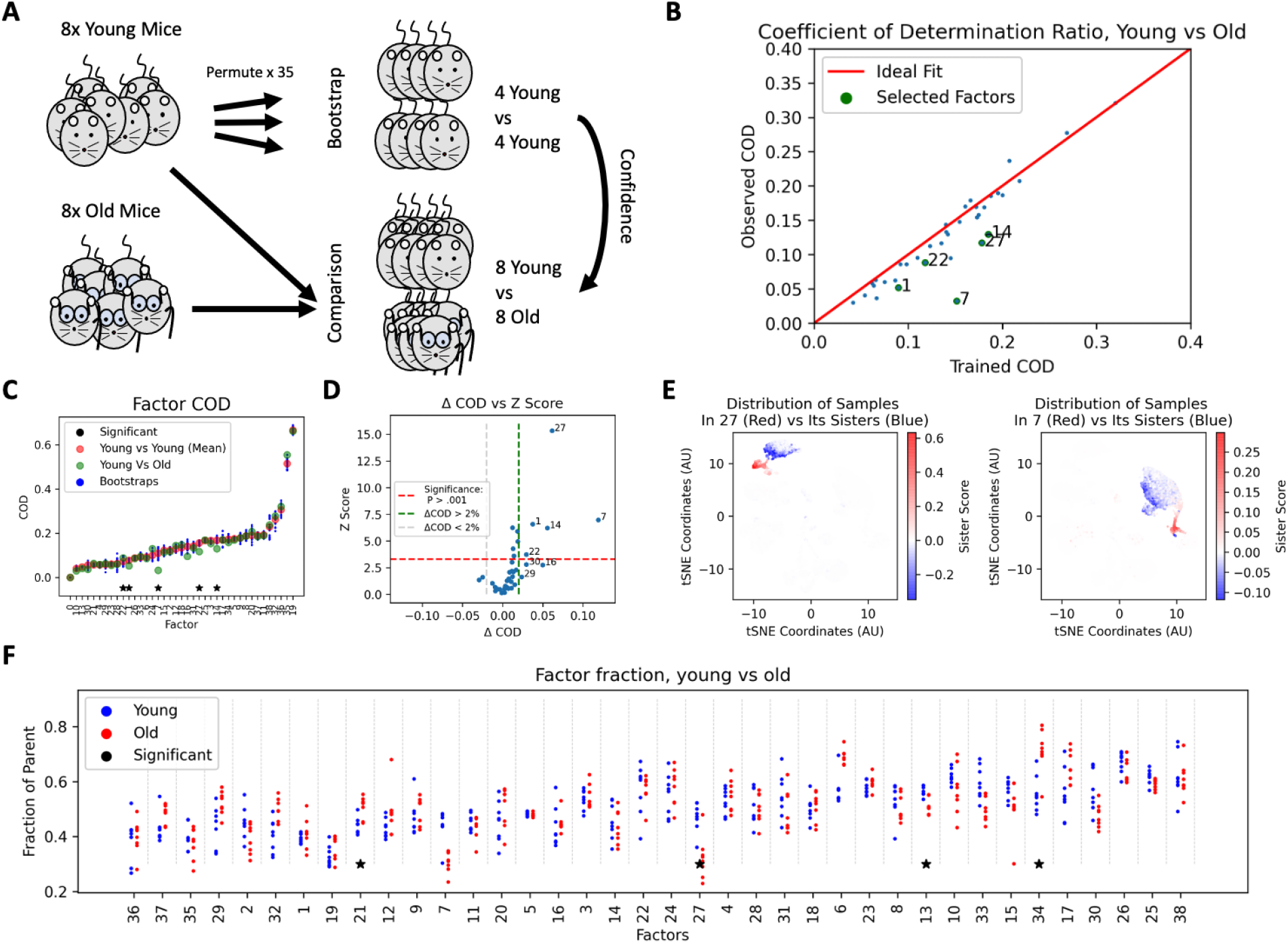
A: Cross-validation scheme. An RFR was trained on all young mice, 35 permutations were established consisting of 4 young mice vs 4 other young mice. The RFR node predictions were calculated using the first 4 young mice and COD was established when predicting the gene expression of the other 4 young mice. B: Mean COD when forest predicts gene expression in young vs old mice for each RFF. RFFs where COD was significantly different in old mice and young mice are labeled. C: COD of each individual factor when trained on 4 young mice and predicting 4 other young mice (blue), mean factor COD across all permutations (red), and COD of each individual factor when trained on 8 young mice and predicting 7 old mice (green). Factors with significant differences in COD are marked with black stars. D: Plot of Z score compared with difference in mean COD in young mice and COD in old mice. Factors where differences are both large and significant are labeled. E: UMAP projection colored by the sister scores for factors 7 and 27, where greatest differences in predictive power lie. F: Mean fraction of the samples captured by nodes in each factor from the parent node is plotted for each mouse, young mice in blue, old mice in red. Factors with significant differences in percent of samples captured between young and old are labeled with black stars.

Conversely, the COD of the NC7 when applied to cells from old mice is only 3.3%, indicating that immature oligodendrocytes in older mice have a significantly different gene expression profile. The expression for test statistics of individual CODs is not known in this context because we are uncertain of the distribution they are drawn from, however assuming them to be approximately t-distributed, 5 factors have significant shifts in prediction quality: NC7, NC27, NC14, NC1, and NC22. (**Fig 5D**)

To establish the biological relevance of this discrepancy in predictive power we can examine the individual genes that have better and worse prediction quality within this cluster. We list the genes that were well predicted by NC7 (COD > 15%) in **Table 2**. Certain key gene expression patterns remain accurately predicted, namely expression of Plp1, Mobp, and Gria2, all key identity markers for immature oligodendrocytes involved in maintaining the physical structure of oligodendrocytes and key physiological functions. Conversely, the predictive quality collapses for almost half of the genes that were previously significantly predicted in this sub-population, namely Fth1, Car2, and Scrg1, which are metabolic proteins responsible for ion metabolism, and Rtn, Tnr, and Bcan which are key transcriptional factors involved in oligodendrocyte differentiation.

**Table 1.** Factor characteristics

| Factor | Identity | Marker Genes | COD Young | COD Old |
| --- | --- | --- | --- | --- |
| 7 | Immature oligo-dendrocytes | Cldn11, Bmp4 | 11% | 3.3% |
| 1 | Klk6 oligo-dendrocytes | Cldn11, Klk6 | 7.7% | 5.2% |
| 27 | Blood immune cells | CD45, Tmem119(-) | 16% | 12% |
| 14 | Monocytes | CD45, Plac8 | 17% | 13% |
| 22<sup>2</sup> | Perivascular Macrophages | CD45, Pf4 | 7.3% | 8.9% |

**Table 2.**
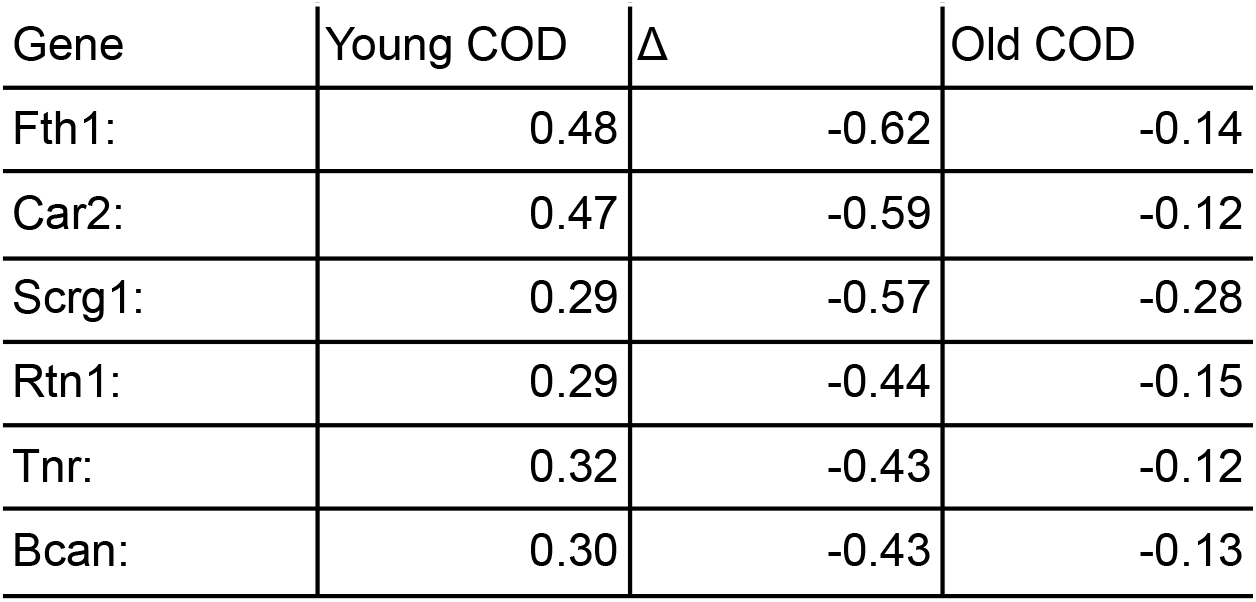
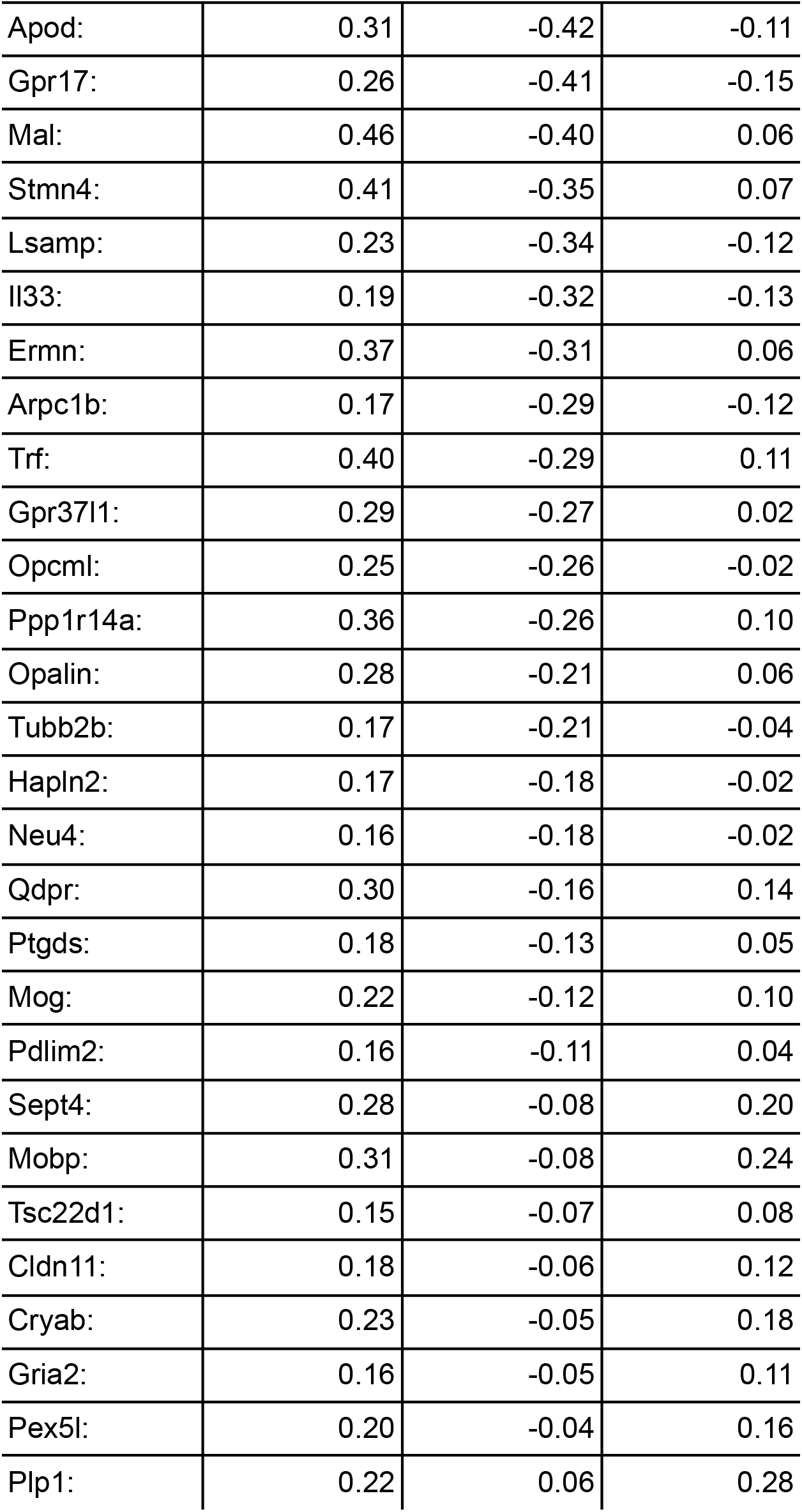
Differentially predicted genes in old vs young, factor 7. Genes are sorted by the difference in the COD among young vs old cells, demonstrating that some key genes retain an almost identical regulatory pattern while others undergo a dramatic change.

We’ve established that we can observe gross changes in expression profiles for certain factors, however we can also use RFFs to track changes in cell populations using our factors. In order to establish the variability of cell populations between different mice, we examined the mean proportion of cells captured from the parent node by nodes in each factor and each mouse (**Fig 5F**). We established the range of percentages of the samples in the parent that are captured by each factor across all 8 young mice and all 8 old mice. To determine whether there was a significant shift in the percentage of the parent samples captured by the nodes in each factor, we performed a Mann-Whitney U Test for the 8 fractions captured in young mice compared to 8 fractions captured in old mice. NC13 exceeds the corrected significance threshold of .00125 (NC13 MWU p=.00097), and several factors reached a significance p<.005 and are marked as well. NC13 is a child to NC3 which contains cells overexpressing Cldn5 (4.3 vs 0.19, t-test p<machine epsilon), which means NC3 contains primarily vascular cells making up the brain blood vessels. NC13 the plurality of sisters for NC13 are in NC21, and Cldn5 is highly expressed in both, so NC13 encodes for a distinction between two subtypes of vascular cells. NC21 predicts over-expression of Vtn, a marker for pericytes, and NC13 predicts over-expression of Cldn5 even relative to NC21, marking them as Endothelial Cells. (Vanlandewijck et al. 2018) The prevalence of NC21 is greater in old cells, indicating that endothelial cells are more prevalent relative to pericytes according to the model postulated by the random forest. Interestingly, the predictive power of NC21 and NC13 only has a small though significant discrepancy (4.7% vs 4.1% COD, T Test, p>.002). In absolute terms the observed changes in populations are small, so attributing significance to them is difficult without a hierarchical model of the type we demonstrate here, which aids in the detection of important effects such as the collapse of immature populations.

## Discussion/Conclusions

The analysis performed by Ximerakis on the mouse brain dataset is performed conventionally, and one of the gold-standard methods used by Ximerakis to analyze the data successfully is firstly to cluster cells coarsely, and then re-cluster the cells in coarse clusters to account for the dramatic changes in behavior that can occur in different parts of datasets. This is a popular technique that works well, as indicated by its wide usage and inclusion in the Seurat example vignettes, but it is a highly manual procedure. A Random Forest Regressor takes a very similar approach by iteratively partitioning the dataset and then treating each partition as an entirely new problem, however RFRs are generally regarded as somewhat opaque tools that don’t provide interpretable information on the structure of the underlying data. The method we’ve described here will help researchers reconsider the manual reclustering procedures that are labor intensive and require judgement calls and the opaque nature of RFRs. We provide a way of gleaning the meaning behind intermediate partitioning steps an RFR, showing us interactions between features that would normally be hidden. In this way, RFR attempts to resolve Simpson’s Paradox by providing a structured way to control

RFRs do come with tradeoffs despite their advantages. RFRs don’t perform as well on highly linear datasets, since each individual estimator in the ensemble ultimately outputs a step function. Additionally, RFs can be overwhelmed by uninformative features if no informative features are present in most bootstraps. Additionally, If feature sparsity is uncorrelated between features and exceeds 50%-70% RFs will generally have trouble selecting split points effectively, since they rely on ranked data. For sparse data we recommend using RF implementations that employ linear combinations of features (either random as originally postulated by Bierman or dimensionally-reduced as postulated by (da Silva, n.d.)). Finally, the number of nodes needed to reflect nested structure doubles for each level of nesting, so if the dataset has a structure that is too deeply nested, the number of nodes that need to be clustered to reflect this structure can become prohibitively large, potentially eliminating the advantages of an RF over network methods or online algorithms like DBSCAN.

Ultimately, the greatest advantages of the RFR factorization method we describe here rest on its unsupervised nature. Researchers can manually glean much of the information we present here by controlling for the right variables or subclustering the right cell populations manually, however if something surprising is hidden in a cell population that is not the primary target of the investigation, it is likely to remain that way unless RFR factorization is employed. Conversely by employing RFR factorization, the researcher will recover more pure and isolated groups of genes corresponding to subtler effects in the data. RFR factorization is easy to tune to provide different levels of detail about the dataset being analyzed, produces intuitive subdivisions, and can be run efficiently even on very large datasets. Our methodology will reveal surprising findings to you when you run it on your single-cell data, and lead you down avenues of investigation that were previously obscured.

## Methods

### RFR Construction

For each RFR, a root node was constructed by partitioning the set of all samples. For each node, if maximum depth or minimum sample quantity was not reached, a split was performed. To perform any split, S samples were bootstrapped with replacement from the set of all samples satisfying the criteria of the node being split and all ancestor nodes forming the subset S’.To perform an unsupervised split, the set of all features M was split randomly into input features and output features I and O. From inputs, F inputs were bootstrapped without replacement forming the subset F’. From outputs G outputs were bootstrapped with replacement forming the subset G’. The S’ x F’ input matrix X’ was constructed by selecting the samples S’ and features F’ from inputs X. The S’ x G’ output matrix Y’ was constructed by selecting the samples S’ and the features G’ from outputs Y. If specified, X’ and Y’ were dimensionally reduced to the top K principal components by NIPALS (Wold 1975), producing a revised input and output pair X’’ and Y’’ with dimension S’ x K and S’ x K. Given the input X’ or X’’ and Y’ or Y’’, for each input feature j (eg each gene) and sample i, potential child node subsets were constructed consisting of subset 1 containing samples where x_i,j_ < x_s,j_ and subset 2 containing samples where x_i,j_ > x_s,j_. The quality of each split was calculated by finding the sum of the squared deviations from the median for each output feature k in the newly formed nodes:

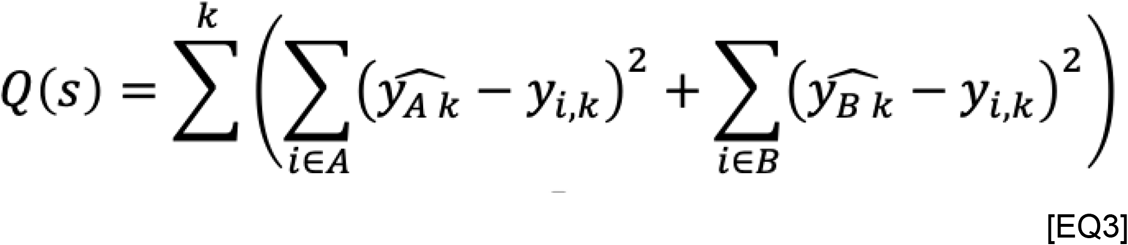

Where A and B are potential children where i is in A or B if x _i,j_ is greater than or less than s_j_ respectively, y-hat_Ak_ is the median of output k in A, y-hat_Bk_ is the median of k in B. After evaluating all possible splits for all selected inputs, the split producing the smallest sum of squared deviations in both child nodes was selected. Two new child nodes were constructed containing all samples that satisfied the conditions selected during the split and the conditions of all ancestor nodes. Alternative error metrics have also been implemented using other robust statistics such as median absolute deviation from the median, sum of absolute deviations from the median, coefficient of determination, and the like.

To predict the gene expression of an unknown sample, the unsupervised forest operates normally, finding the nodes whose split criteria match the sample and then taking the conditional estimates of those nodes and summing them. This is mathematically equivalent to taking the mean of the predictions of all leaves to which the sample is assigned. At face value this may imply that when acting as a regressor the forest allows features to predict their own behavior, however this does not reflect a dangerous practice because features are prevented from self-predicting during forest training.

Procedure was implemented in Rust, and all code including code used in the generation of figures is available on github at https://github.com/bbrener1/rf_5

### Node Representation Construction

Node representations were constructed by obtaining all nodes comprising all trees in an RFR in left-to-right order and for each node computing one of three summaries. To construct a Node-Sample Encoding each node was summarized by an array of length N, where each boolean was true if a sample was present in the node and false otherwise. To construct a Node-Sister Encoding each node was summarized with an array of of length N where the ith integer was 1 if sample i was present in the node, -1 if sample i was present in the sister node, and 0 otherwise. To construct a Node-Gain Encoding, we first found the mean for each output feature among the samples present in the node and the samples present in the parent of the node. The summary of a node was constructed by finding the difference between the parent output means and the node output means. For each encoding, the node summaries were arranged into a rectangular matrix Nd x N or Nd x G in dimension where Nd is the total number of nodes present in a Forest.

### Node Cluster Construction

Node clusters were inferred by first constructing a k-nearest-neighbor graph of node summaries using the specified metric (cosine distance by default), with a default k parameter of 100. After constructing the neighbor graph, the python package Community was used to partition the nodes by graph connectivity using default parameters. Briefly, Community uses a graph-theoretic approach to partitioning samples on the basis of a network connecting individual samples to each other using sequential cuts to the connecting edges and observing the number of cuts required to produce isolated graphs. By constructing a k-nearest-neighbors graph on a dataset using a distance metric of choice, community detection can be performed on numeric data. Community was selected as a clustering approach because it is well behaved when intra-cluster distance variance is inconsistent between different clusters, rendering it superior to DBSCAN and other conventional methods.

### Mouse Brain Dataset

Refer to Ximerakis for full specification. Brains of 8 young mice and 8 old mice were harvested and homogenized into a single-cell suspension, tagged with 10X chemistry and sequenced to saturation. The resulting dataset contained 16027 cells originating from young mice and 21041 cells from old mice after filtering for quality. Alignment and quantification was performed by Cell Ranger, cells were log-normalized, repooled, and scaled.

## Acknowledgements

This work was supported, in part, by National Institutes of Health award R24DK106766-01A1 to M.C.S. A.B. was supported by NIGMS R35GM139580

We wish to thank James Taylor for his contributions to this paper, he nurtured its most foundational ideas and was instrumental in its publication; we are deeply saddened that he could not be with us to see its release.

## Abbreviations

If abbreviations are used in the text they should be defined in the text at first use.

scRNAseq: Single-cell RNA sequencing
RF: random forest
RFR: random forest regressor (to distinguish from a random forest classifier)

## Supplemental Information

Default RFR parameters:

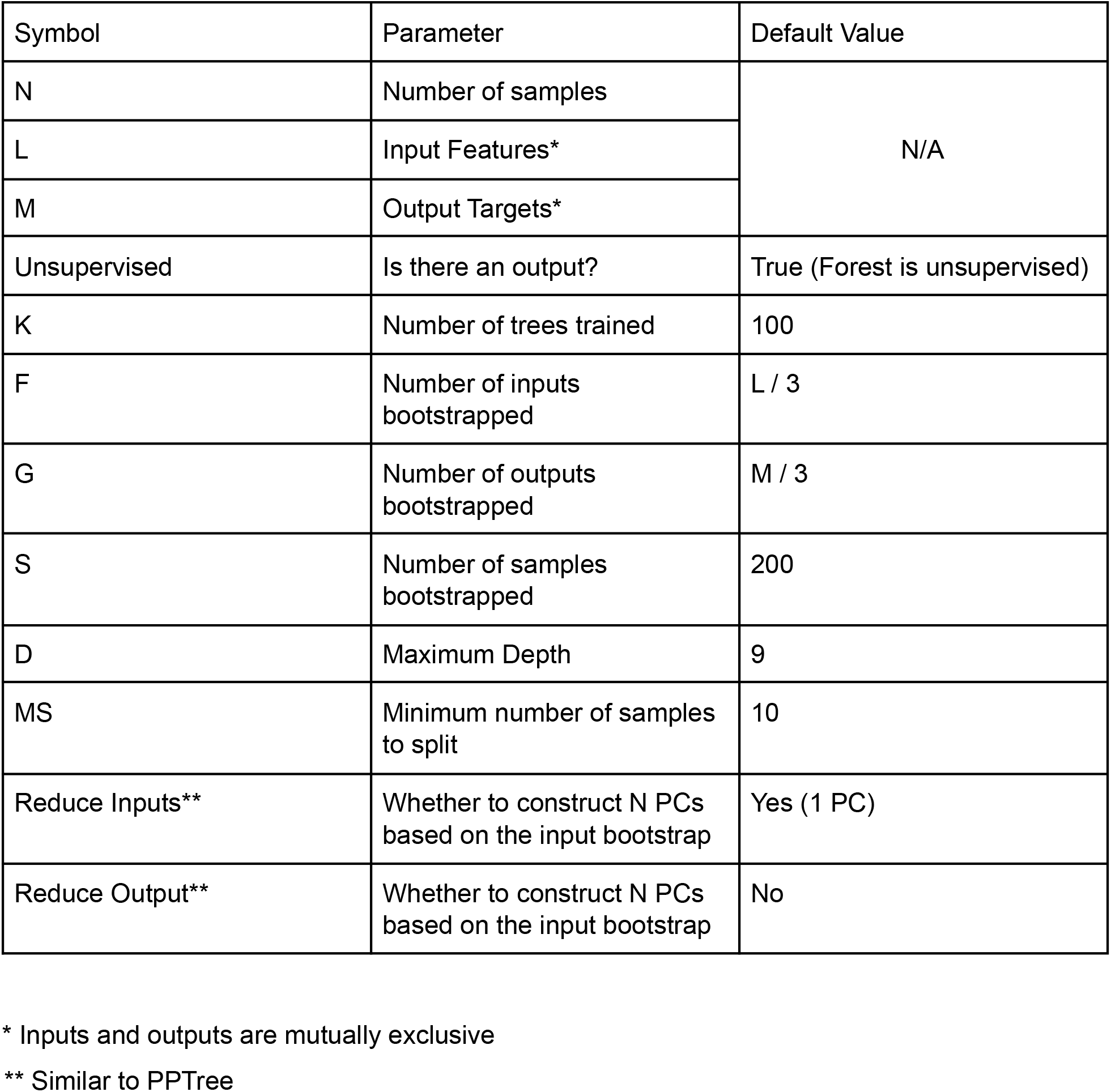

| Symbol | Parameter | Default Value |
| --- | --- | --- |
| N | Number of samples | N/A |
| L | Input Features* |  |
| M | Output Targets* |  |
| Unsupervised | Is there an output? | True (Forest is unsupervised) |
| K | Number of trees trained | 100 |
| F | Number of inputs bootstrapped | L / 3 |
| G | Number of outputs bootstrapped | M / 3 |
| S | Number of samples bootstrapped | 200 |
| D | Maximum Depth | 9 |
| MS | Minimum number of samples to split | 10 |
| Reduce Inputs** | Whether to construct N PCs based on the input bootstrap | Yes (1 PC) |
| Reduce Output** | Whether to construct N PCs based on the input bootstrap | No |

\* Inputs and outputs are mutually exclusive
\*\* Similar to PPTree

scRNAseq - Single Cell RNA Sequencing

RFR - Random Forest Regressor

## Supplemental Figures

**Supplemental Figure 1:**
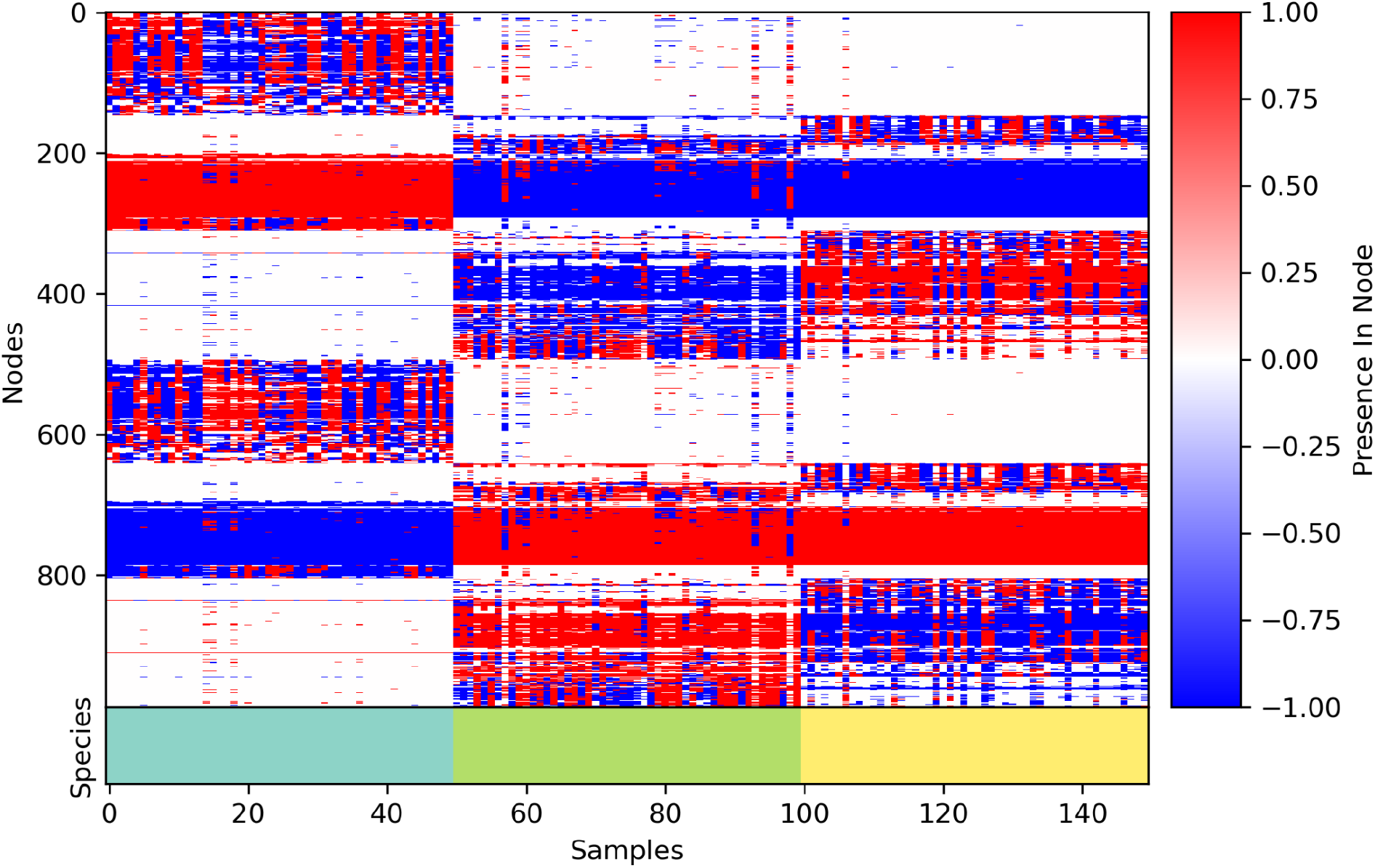
Iris dataset node-sister encoding vs true iris species. We trained an unsupervised forest on the iris dataset without including iris species in the training data and plotted the node-sister encoding. The true iris species is labeled by color at the bottom, demonstrating that the forest has a hierarchical structure that can distinguish all three Iris species, while recognizing the similarity of two clusters to each other as being greater than the similarity of the third cluster.

**Supplemental Figure 2A:**
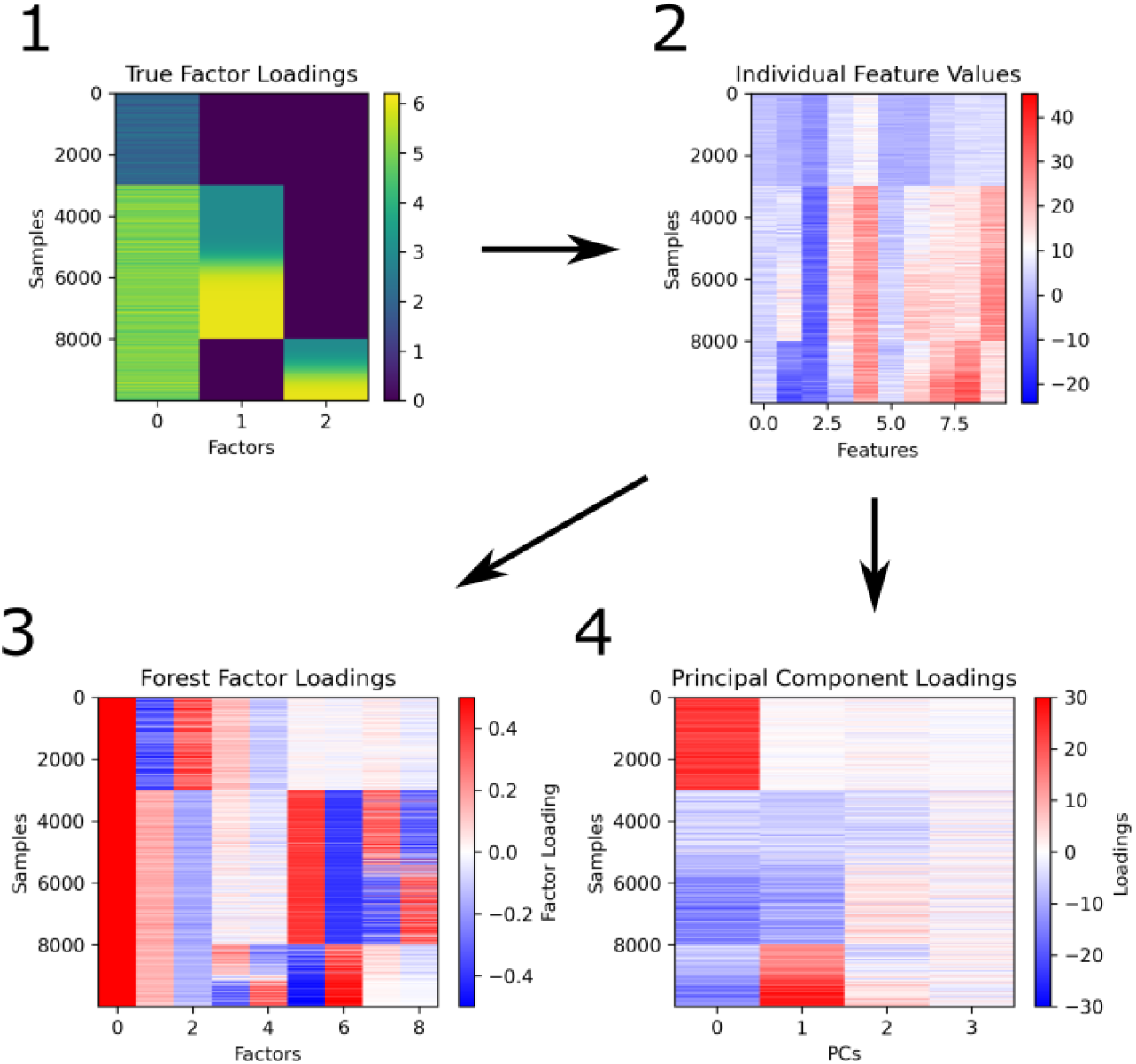
RFR reconstruction of latent factors. Three latent factor loadings were constructed (A1), and used to generate 10000 samples with 10 simulated features (A2). The simulated data in A2 was analyzed by forest factorization (A3) and PCA (A4) to obtain the inferred factor loadings.

**Supplemental Figure 2B:**
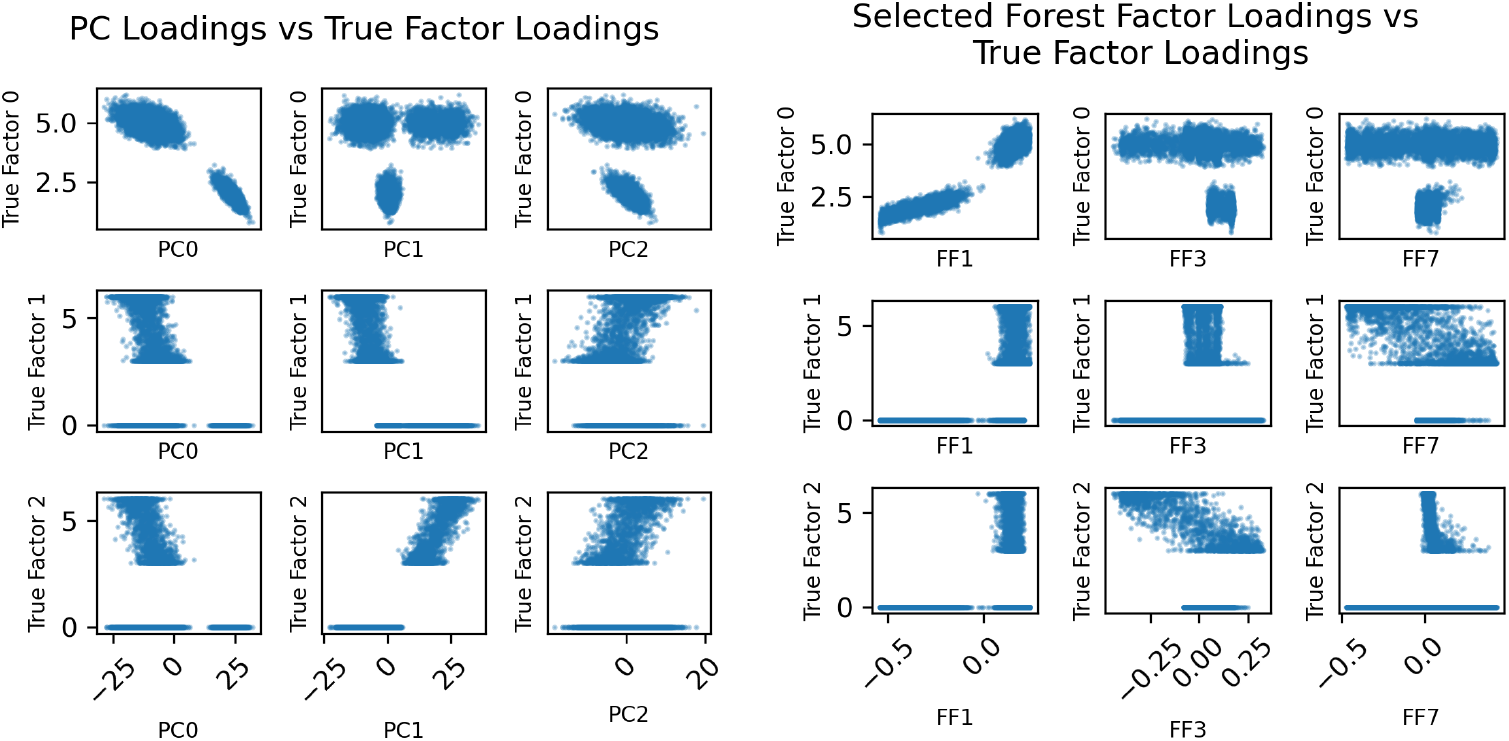
Correlations of the inferred PC loadings and forest factor loadings (sister scores) to true simulated loadings. We plot PC loadings of the top 3 PCs and the sister scores of selected forest factors against the true loadings of the simulated factors to observe how accurate the recovery of the true values is. Note that the loadings of PC1 and PC2 both have some correlation to the loadings of true factors 1 and 2. PCs are unable to disentangle these naively correlated effects. In comparison, note that forest factors 7 and 3 are strongly correlated to true factors 1 and 2 respectively, but NOT correlated to the other. This shows that forest factorisation can disentangle non-linear dependency in a way which is difficult for PCA.

## Footnotes

1 NC7 is not the only factor coding for Cldn11-positive oligodendrocytes, so there is appreciable expression in “all other cells”, the factor coding for all Cldn11-positive oligodendrocytes is NC19, for which the statistic is 3.9 TMP vs 0.094 TPM, p<machine epsilon

2 Note the increase in COD in older cells. It is unclear why COD would increase in older cells, though potentially a rare subpopulation that was co-clustering with cells in 14 is no longer present

## Notes

https://github.com/bbrener1/rusty_axe

### Competing Interest Statement

The authors have declared no competing interest.

